# Lessons from combined metabolic model of mesophyll and guard cells

**DOI:** 10.1101/2024.05.30.596642

**Authors:** Devlina Sarkar, Sudip Kundu

## Abstract

Understanding of overall metabolisms of guard cell (GC) and mesophyll cell (MC), their possible interactions and roles in stomatal movement will help in designing crop plants with higher water use efficiencies. Although, previous constraint based modelling and analyses correctly predicted some known metabolic patterns of isolated C_3_ MC, GC and Crassulacean Acid Metabolism (CAM) MC, quantitative estimations of combined metabolism of GC and MC and detailed analysis of GC metabolism in CAM are still awaiting. A six phase diel combined model of GC and MC is constructed integrating existing models of two different cell types with necessary modifications guided by known physiology. It is used to predict the similarities and dissimilarities of GC and MC metabolisms. In addition to previously predicted results, analysis of this new two-cell model successfully shows higher activities of some experimentally observed C_4_ like enzymes in GC than MC in C_3_, the possible storage patterns of the osmolytes like K^+^, malate^2-^, sucrose etc., in CAM GC. Results also show different patterns of starch synthesis and accumulation, ATP production and utilization in GC and MC of C_3_ and CAM. This combined model integrates stomatal opening, gaseous exchange and GC-MC metabolisms. It is a significant step towards understanding and quantitative estimation of the whole leaf tissue metabolism linking gaseous exchange with environment.

**Significance statement:** Our time-resolved combined metabolic model of guard and mesophyll cells integrates stomatal opening, gaseous exchange and cellular metabolisms in C_3_, CAM and partial CAM conditions; their analyses provide quantitative estimations of metabolic fluxes, ATP production and utilization, and different metabolic patterns of starch synthesis and accumulation of both the cells. It is a significant step towards understanding and quantitative estimation of the whole leaf tissue metabolism linking gaseous exchange with environment and phloem loading.

## 1. Introduction

Rapid increase in the world’s population, continuous decrease in available water, and arid environment will surely impact food security in near future (Galanakis, 2024). To face this challenge, plant biotechnologists are trying to integrate Crassulacean Acid Metabolism (CAM) into crop varieties. Success of this endeavour demands understanding of many different aspects of CAM and non CAM plants including their anatomy and physiology. The gaseous exchange between the environment and mesophyll cells (MCs) of plant system is regulated by microscopic pores, called stomata, present in the epidermal layer of plants. The dynamic control of stomatal aperture is facilitated by two specialized cells known as guard cells (GCs). In C_3_, stomata are open at day, but in CAM, the scenario is opposite to prevent the water loss through transpiration at daytime. Various ions and osmolytes play pivotal role, in opening of stomata, facilitating gaseous exchange between environment and internal leaf airspace in both C_3_ and CAM. In C_3_ plants, the ATPase-Proton pump is activated during the daytime in response to light, leading to the extrusion of protons from the GC (Shimazaki *et al*., 2007). In response to the hyperpolarization across the membrane, potassium ions (K^+^) enter the GC to maintain charge balance. The positive charge is primarily offset by malate^2-^ produced from starch breakdown and chloride ions (Cl^-^) uptake. Additionally, sucrose generated from starch breakdown in MC is transported to GC through the apoplastic region (Lawson, 2009). The elevated concentrations of these metabolites reduce the water potential inside the GC, thereby increasing osmotic pressure (OP) and prompting the water to enter the GC from the surrounding environment via osmosis. This influx of water enhances the turgidity of the GC, leading to the opening of the stomatal pore (Lawson, 2009; Lawson *et al*., 2014). In C_3_ plants, experimental evidence suggests that K^+^ accumulates rapidly in the early morning, while sucrose primarily balances OP in the afternoon (Talbott and Zeiger, 1996). However, the mechanism underlying stomatal opening at night in CAM plants remains a topic of investigation. One hypothesis proposed by Lee, 2010, suggests that H^+^ pumps are activated during darkness, leading to the entry of K^+^ into the GC.

In C_3_ plants, sucrose accumulation during the latter part of the daytime, plays a role in regulating OP and the opening of stomata. Although, important role of sucrose is also known in CAM plants (Lee, 2010), the precise timing of its accumulation during the night to maintain OP in GC and its influence on gluconeogenesis are still not fully understood. A comprehensive analysis of charge accumulation, interactions of GC and MC, and how these are related with stomatal behaviour is needed.

Moreover, among the vast variety of CAM species, CAM cycling and CAM idling are considered to be the precursors of CAM, representing intermediary stages in the evolutionary transition from C_3_ to CAM plants (Cushman and Bohnert, 1997). In CAM cycling, stomata open during the day and close at night, whereas in CAM idling, there is no gaseous exchange. In recent years, researchers have been endeavouring to introduce CAM traits into C_3_ plants through gene editing and synthetic biology techniques. Incorporating CAM traits into C_3_ requires an understanding of the enzymatic and regulatory pathways of C_3_ and CAM photosynthesis at system level (Yang *et al*., 2015; Borland *et al*., 2014; Yuan *et al*., 2020). Constraint based modelling approach was used to study CAM MC photosynthesis extensively (Cheung *et al*., 2014). Recent studies have demonstrated that introducing CAM into C_3_ plants can result in crop varieties with improved water use efficiency (Shameer *et al*., 2018; Töpfer *et al*., 2020). These studies focus on modification of metabolism of MC to introduce CAM in C_3_ plants. Whereas, besides MC, modifications of stomatal behaviour and hence metabolism of GC is also important for improving photosynthetic and water use efficiency of C_3_ plants in arid environments (Nunes-Nesi *et al*., 2007; Daloso *et al*., 2016b). Some previous experimental and theoretical studies explored the behaviour and metabolism of isolated GCs in C_3_ plants (Lawson and Matthews, 2020; Tan and Cheung, 2020; Lima *et al*., 2023). Robaina-Estévez et al., 2017 showed the differences in metabolic fluxes between the isolated MC and GC of *Arabidopsis thaliana,* mainly focusing on central carbon metabolism of GC and revealed the role of the tricarboxylic acid (TCA) cycle in malate synthesis and the Calvin–Benson cycle in sucrose synthesis. Tan and Cheung, 2020 demonstrated the metabolism of isolated C_3_ GC and how the balance of charge and accumulation of osmolytes are maintained to regulate OP. Previous experimental studies reported the role of anions in CAM GC (Kong *et al*., 2020; Lefoulon *et al*., 2020), transformation from C_3_ to CAM metabolism in common ice plant under draught or salt stress (Kong *et al*., 2020; Qiu *et al*., 2023), and differential activities of some enzymes in GC and MC, but in isolated protoplasts (Zhu *et al*., 2009; Reckmann *et al*., 1990; Bates *et al*., 2012). However, experimental studies have revealed variations in the response of isolated stomata compared to those surrounded by MC when exposed to red light (Lee and Bowling, 1992). Similarly, mesophyll cells may influence guard cell metabolism as well. Therefore, a detailed understanding of interactions of GC and MC is needed to understand their integrated metabolism and the influence of MC in stomatal regulation in both C_3_ and CAM conditions.

Here, we constructed a six-phase diel metabolic model, comprising both GC and MC, to capture their metabolisms and interactions. We link the osmotic pressure (OP) of GC to the gaseous exchange between environment and internal airspace within leaf mimicking the dependency of stomatal opening on osmolyte accumulation in GC. Simulations successfully predict the known characteristics of GC and MC of C_3_ and CAM plants. Results show higher activities of some enzymes like phospoenolpyruvate carboxylase (PEPc), NAD(P)-malate dehydrogenase (MDH), malic Enzyme (ME), pyruvate, orthodiphosphate dikinase (PPDK), fumarase, succinate dehydrogenase (SDH), carbonic anhydrase (CA) etc., in GC than MC in C_3_ plant. Expansion of our study to simulate CAM conditions provides insight into the collaborative functioning of GC and MC in CAM plants, as well as the metabolic and energetic variations of GC and MC in C_3_, CAM and partial CAM conditions.

## 2. Results and discussion

### 2.1 Construction of a six-phase diel metabolic model comprising a mesophyll cell and a guard cell

A six-phase diel combined model of a GC and a MC is developed using the mass and charged balanced core leaf metabolic model of Töpfer et al., 2020. Firstly, this core leaf cell model is duplicated six times and storage reactions of different ions and organic acids are added to construct a six-phase model of a single cell. Then we duplicate the cell to represent each as GC and MC. GC and MC are linked by sucrose transport from MC to GC. The six phases are named as phase 1 to phase 6 to represent dawn, mid-day, afternoon, dusk, night and end of night respectively. The total time of 24 hours of a complete diel cycle is distributed in these six phases in a ratio of 1:10:1:1:10:1. The storage reactions are added in such a way that metabolites stored in each phase are transferred to next phase. In GC, there is no production of phloem sap, making MC the only source of phloem sap production. Size of the vacuole is not constrained. We have integrated the metabolisms of GC and MC by linking the increment of OP with CO_2_ availability to MC. Higher the OP, higher is the available CO_2_ to MC. Ions (K^+^, Cl^-^, malate^-2^) are considered to be accumulated in GC to maintain OP. As malate plays a more significant role than Cl^-^ (Lee, 2010) and to simplify the interpretations of model predictions (Tan and Cheung, 2020), we have allowed the accumulation of only malate to balance the positive charge of K^+^ in our model and termed it as the “standard scenario”. We have also analysed our model by allowing the accumulation of only Cl^-^ and both malate and Cl^-^ together (described later in subsection 2.8). Henceforth, we have presented results of simulation of standard scenario unless it is specifically mentioned. The model files can be found in the Data S1 and the model is represented schematically in Figure 1. The details of methods are described later in the “Experimental procedure” section and the general constraints used on metabolisms of GC and MC in C_3_, CAM and partial CAM conditions are summarized in Table 1.

**Figure 1.**
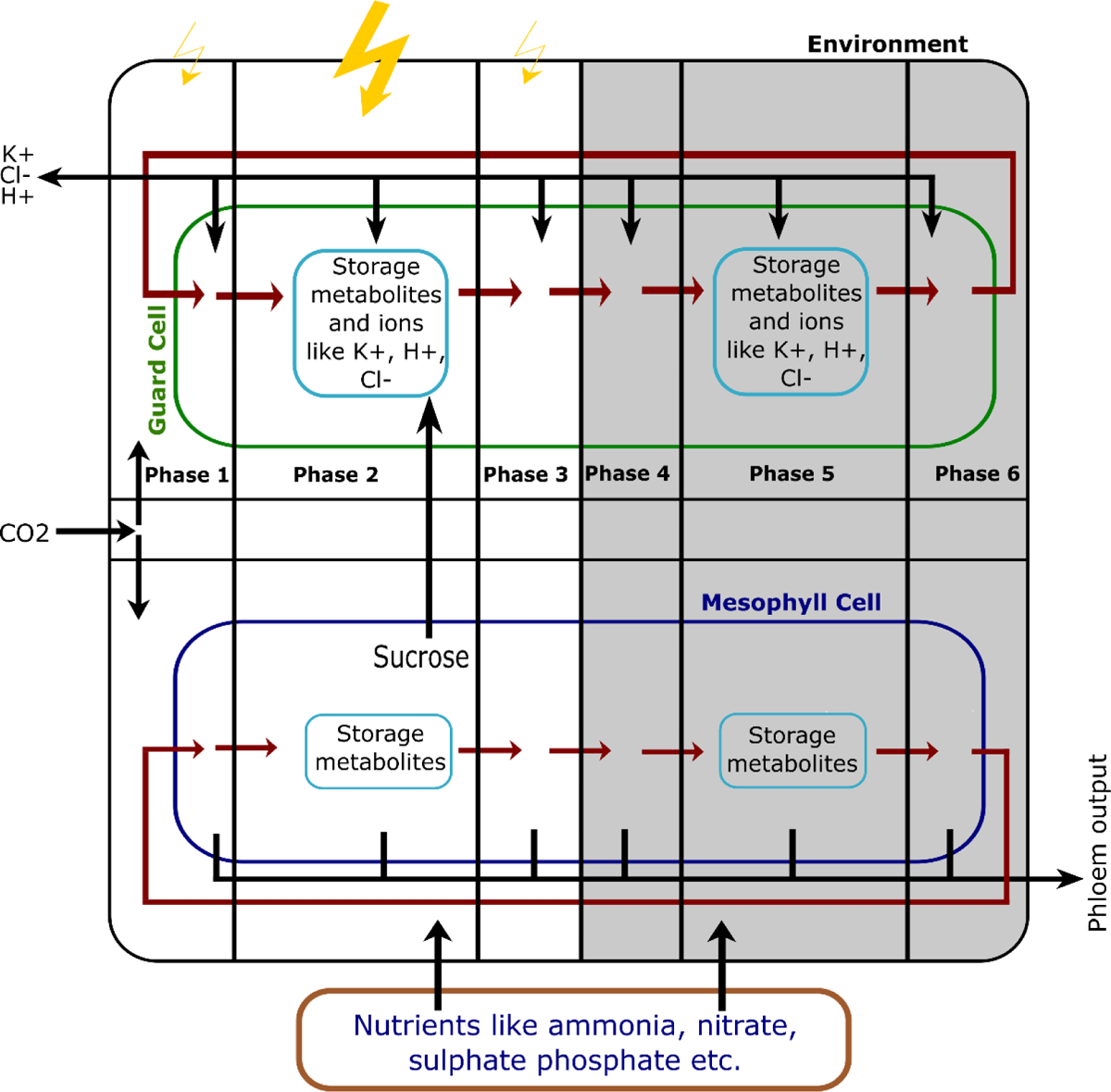
Schematic representation of our six phase diel combined model of mesophyll and guard cells. Light is available to the first three phases. CO_2_ entry from environment is linked to OP in GC and depending on the constraint on OP, CO_2_ is made available in all the phases (not shown here). Exchange of O_2_ has also been limited depending on the OP. Nutrients are available to all the phases of day and night. Sucrose transport from MC to GC is shown here for C_3_ and CAM cycling conditions. For CAM and CAM idling, it is transferred in the phase 4 (not shown here). All other constraints are summarized in the table 1.

**Table 1.** Constraints applied to GC and MC in C_3_, CAM, CAM cycling and CAM idling.

| Type of Cell | Constraints | Values |
| --- | --- | --- |
| MC | Photon ratio in the three phases of day | 1:10:1 |
|  | Nitrate uptake ratio at day and night* | 3:2 |
| | Phloem output rate (in $\mu\text{mol.m}^{-2}.\text{s}^{-1}$) | 0.259 |
|  | Phloem output ratio at day and night** | 3:1 |
|  | % of total accumulated osmolyte in GC to be sucrose,<br>transferred from MC to GC in phase 2 in C<sub>3</sub> and CAM cycling | 80 |
|  | % of total accumulated osmolyte in GC to be sucrose,<br>transferred from MC to GC in phase 4 in CAM and CAM idling | 80 |
|  | C:O of RuBisCO in each phase of day in C<sub>3</sub> and CAM cycling | 3:1 |
|  | C:O of RuBisCO in each phase of day in CAM and CAM idling | 5.15:1 |
| GC | Photon ratio in the three phases of day | 1:10:1 |
|  | Nitrate uptake | 0 |
|  | Phloem output | 0 |
|  | C:O of RuBisCO in each phase of day in C<sub>3</sub> and CAM cycling | 3:1 |
|  | C:O of RuBisCO in each phase of day in CAM and CAM idling | 5.15:1 |

\*Further distributed according to the length of the phases
\*\*Free to be produced in any phase of day and night

### 2.2 Model successfully predicts the known facts and differences in metabolisms and enzymatic activities in GC and MC

Our combined metabolic model of GC and MC helps us to integrate (i) gaseous exchange through stomatal opening with increase in osmolyte concentration in GC and (ii) transport of metabolites from MC to GC, and (iii) also GC and MC metabolisms with phloem sap production. While previous experimental reports helped us to fix phase and cell specific constraints, simulations provide the relative flux distributions along different phases of GC and MC in all four different metabolic varieties-C_3_, CAM, CAM cycling and CAM idling. The flux distributions (Data S2) in general match with previously predicted metabolic patterns and energetics, observed in analyses of previous models of isolated GC of C_3_ (Robaina-Estévez et al., 2017; Tan and Cheung, 2020) and MC of all four variants (Shameer et al., 2018; Tay et al., 2021), as well as provide a few new insights (Table S1). However, there is no previous study on analyzing the ATP production and utilization of GC in C_3_ and CAM. Our six-phase diel combined model accurately predicts several experimentally observed traits of both GC and MC in C_3_ and CAM plants (Figure 2a,b), as described in the following subsections. Moreover, the predicted flux distributions are able to capture a large number of experimental observations suggesting differential enzymatic activities of GC and MC in C_3_ metabolism (Table 2, Figure 3a,b). A detailed analysis also suggests different activities of enzymes involved in starch synthesis and TCA cycle in different phases in GC and MC of C_3_ and CAM (Figure 4a,b,c,d,e,f) and a detailed account of energy balance and the participating reactions in these cells (Figure 5a,b,c,d).

**Figure 2.**
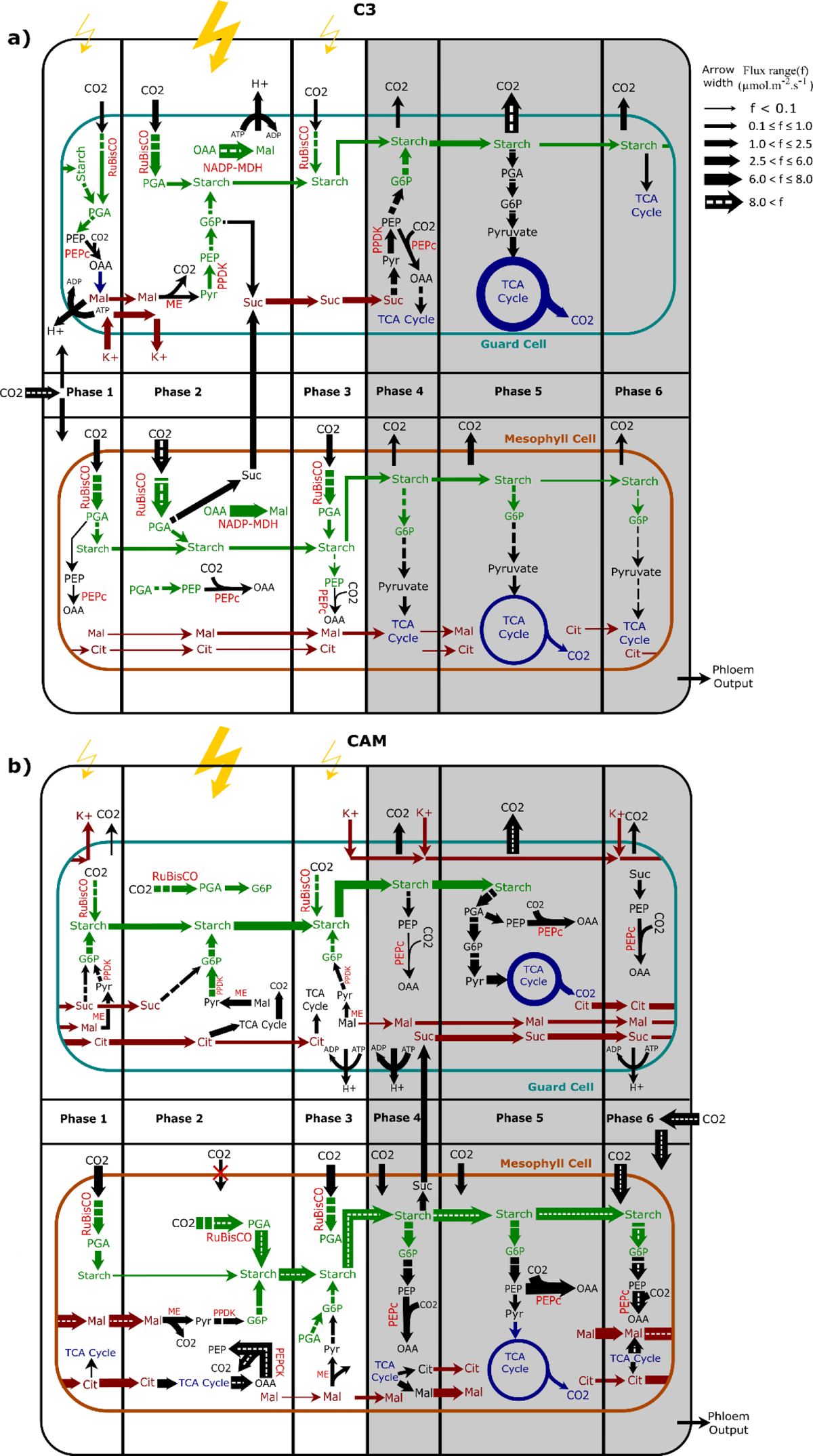
Flux distribution map of accumulation of different osmolytes in GC and metabolic pathways in GC and MC in different phases of a) C_3_ and b) CAM. Different widths of arrows show relative magnitudes of fluxes through the reactions scaled throughout the whole figure. The broken arrows represent a series reactions between two metabolites and dotted arrows represent fluxes of large magnitude that could not be scaled. Different colours represent reactions of different compartments. Green: chloroplast, blue: mitochondria, brown: vacuole and black: cytosol and other general transport reactions. Yellow symbol represents the photon flux to corresponding phase and these arrow widths are not scaled relative to other arrows in the figure. Unit of flux: µmol.m^-2^.s^-1^. Enzymes are coloured in red. ME, malic enzyme; RuBisCO, ribulose-1,5-bisphosphate carboxylase/oxygenase; PEPc, phospoenolpyruvate carboxylase; PEPCK, phospoenolpyruvate carboxykinase; PPDK, pyruvate, orthodiphosphate dikinase; NADP-MDH, NADP-malate dehydrogenase; pyr, pyruvate; PEP, phospoenolpyruvate; OAA, oxaloacetic acid; Mal, malate; Cit, citrate; PGA, 3-phosphoglyceraldehyde; G6P, glucose-6-phosphate; Suc, sucrose.

**Figure 3.**
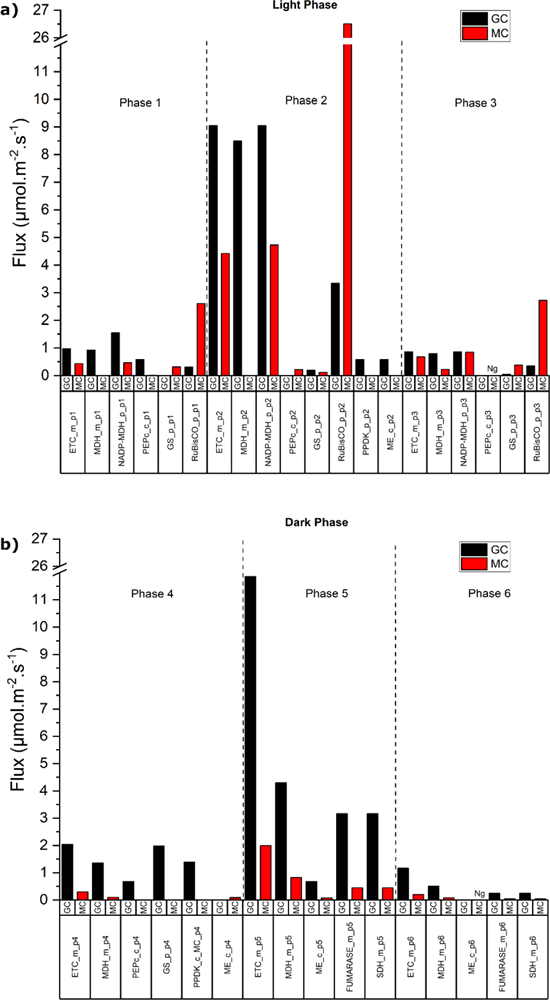
Bar graphs represent relative fluxes through different reactions in mesophyll and guard cellse of C_3_ metabolism in a) light phase and b) dark phase. Black and red bars represent fluxes through the reactions of GC and MC respectively. “_c”, “_p” and “_m” represent cytosol, plastid and mitochondria respectively and p1 to p6 represent the six phases. Ng represents negligible fluxes through the reactions. Unit of flux: µmol.m^-2^.s^-1^. ME, malic enzyme; RuBisCO, ribulose-1,5-bisphosphate carboxylase/oxygenase; PEPc, phospoenolpyruvate carboxylase; PEPCK, phospoenolpyruvate carboxykinase; PPDK, pyruvate, orthodiphosphate dikinase; NADP-MDH, NADP-malate dehydrogenase; GS, glycogen synthase; MDH, malate dehydrogenase; ETC, electron transport chain; SDH, succinate dehydrogenase.

**Figure 4.**
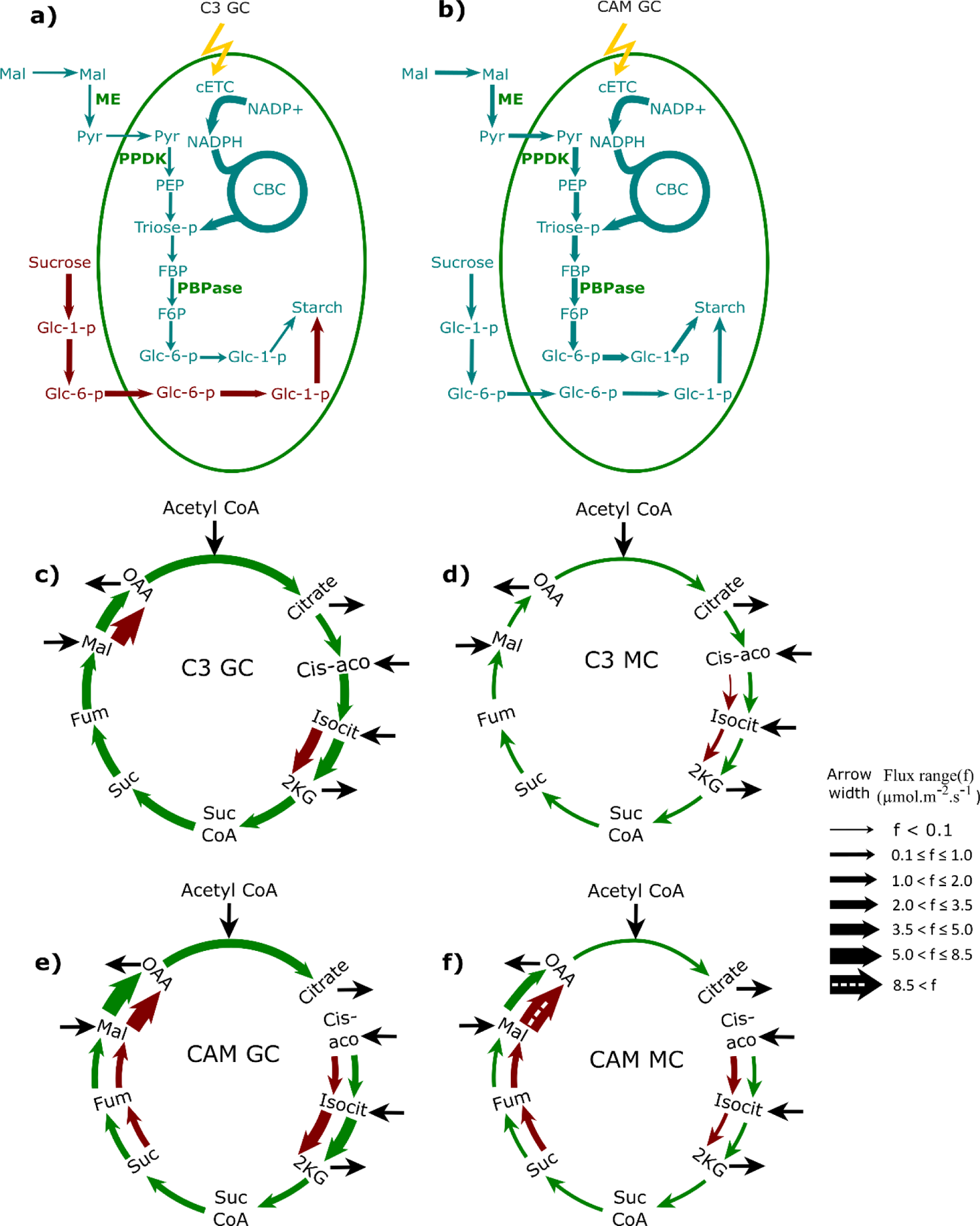
Starch synthesis and activity of TCA cycle during day and night. Starch synthesis pattern and activity of FBPase and PPDK at day (teal blue colour) and night (brown colour) are shown in a) C_3_ and b) CAM. Different modes and flux range of TCA cycle in phase 2 (brown colour) and phase 5 (green colour) are shown in c) C_3_ GC, d) C_3_ MC, e) CAM GC and f) CAM MC. Widths of the arrows represent relative magnitudes of the fluxes through the reactions. The Dotted arrow represents excessive flux that could not be scaled in the diagram. The horizontal black arrows show the transporter reactions of the metabolites between cytosol and mitochondria and the vertical black arrow represents involvement of the metabolite in that reaction. Widths of black arrows do not represent the magnitudes of flux.

**Figure 5.**
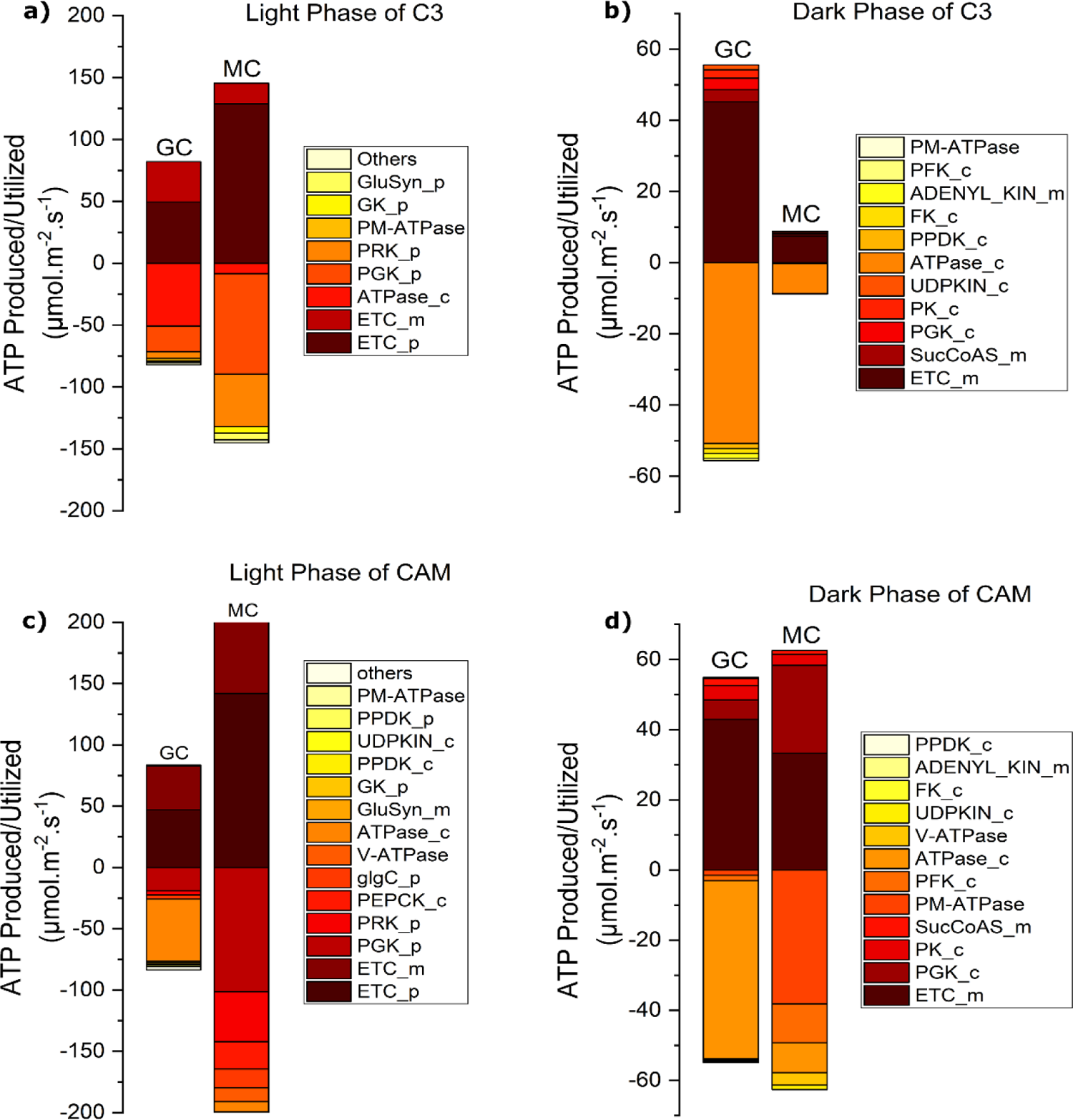
Production and utilization of ATP in GC and MC in light and dark phases of C_3_ and CAM. The bar graphs show major reactions involved in ATP production and utilization in GC and MC of a) C_3_ during light phase, b) C_3_ during dark phase, c) CAM during light phase, and d) CAM during dark phase. The positive and negative values of flux represent the production and consumption of ATP respectively. Unit of flux: µmol.m^-2^.s^-1^. “_c”, “_p” and “_m” represent cytosol, plastid and mitochondria respectively. PEPCK, phospoenolpyruvate carboxykinase; PPDK, pyruvate, orthodiphosphate dikinase; ETC, electron transport chain; GluSyn, glutamine synthase; GK, glycerate 3-kinase; PM-ATPase, plasma membrane proton pump; V-ATPase, vacuolar proton pump; PFK, phosphofructokinase; PGK, phosphoglycerate kinase; SucCoAS, succinyl-CoA synthase; PK, pyruvate kinase; PRK, phosphoribulokinase; FK, fructokinase; Adenyl_Kin, adenylate kinase; UDPkin, UDP Kinase; glgC, glucose-1-phosphate adenylyltransferase.

**Table 2.** Relative flux relation of enzymes in guard and mesophyll cells in C_3_.

| Enzymes | EC number | Relative flux relation | References |
| --- | --- | --- | --- |
| RuBisCO at day | 4.1.1.39 | GC < MC | (Reckmann *et al.*, 1990; Shimazaki and Okayama, 1990) |
| PEPc at day and night | 4.1.1.31 | GC > MC | (Bauer *et al.*, 2013; Leonhardt *et al.*, 2004; Vani and Raghavendra, 1994) |
| NADP-MDH at day | 1.1.1.82 | GC > MC | (Shimazaki2, 1989; Gotow *et al.*, 1985) |
| MDH at day and night | 1.1.1.37 | GC > MC | (Vani and Raghavendra, 1994; Gotow *et al.*, 1985; Leonhardt *et al.*, 2004; Wang *et al.*, 2011) |
| ME at day and night | 1.1.1.40 | GC > MC | (Outlaw *et al.*, 1981) |
| PPDK at day and night | 2.7.9.1 | GC > MC | (Daloso, dos Anjos, *et al.*, 2016) |
| Fumarase at night | 4.2.1.2 | GC > MC | (Vani and Raghavendra, 1994) |
| SDH at night | 1.3.5.1 | GC > MC | (Vani and Raghavendra, 1994) |
| GS at day | 2.4.1.21 | GC < MC | (Robinson and Preiss, 1987) |
| CA at day and night | 4.2.1.1 | GC > MC | (Leonhardt *et al.*, 2004) |

### 2.3 Accumulation of osmolytes, organic acids and activities of proton pumps

Our six-phase model successfully predicts the proton efflux, influx of K^+^, and starch synthesis and breakdown in GC in standard scenario. We restricted the transfer of K^+^ from day to night as the accumulation of K^+^ is experimentally reported to be at daytime in C_3_. We observe its accumulation mainly in the early phases, with OP being maintained during the mid-day and afternoon phases by sucrose produced in GC and imported from MC. These results are in accordance with experimental observation (Talbott and Zeiger, 1996). To balance the positive charge of K^+^, accumulation of malate is observed in GC during the first phase of the day. Whereas in MC, there is a minimal accumulation of malate during all the phases of the daytime. A little nocturnal accumulation of citrate is also observed in MC.

Conversely, in CAM, there is no constraint on K^+^ accumulation. Results indicate that K^+^ starts entering GC in phase 3 and an increase in K^+^ level is observed during the last phase of the night. In phase 3, there is no accumulation of sucrose is observed and OP is maintained by the accumulation of K^+^ only. To balance this positive charge of K^+^ in GC, a small amount of malate is also observed to accumulate during the daytime, whereas it increases at night, along with the increased accumulation of K^+^. During the night phases, both malate and sucrose, along with increased K^+^, contribute to maintain the osmolyte concentration in GC for keeping stomata open. However, at dawn, there is an efflux of K^+^ with OP being sustained solely by sucrose. Furthermore, proton efflux starts in phase 3, which amplifies during the night, aligning with the complete opening of stomatal pores. Reports on CAM GC provide additional support to our findings, suggesting that proton pump activation occurs during darkness, alongside the nocturnal entry of K^+^ into GC (Dayanandan and Kaufman, 1975). To validate these results, further experimental studies focusing on CAM GC are needed.

### 2.4 Different patterns of starch production and utilization in GC and MC of C_3_ and CAM

In C_3_ GC (Figure 2a, 3a,b), starch accumulation initiates post-dawn and persists throughout the daytime, peaking during the 1^st^ phase of night. In phase 2 and 3, CO_2_ is fixed by the enzyme RuBisCO and the produced triose sugar is utilized in producing starch. Whereas in phase 4, dark CO_2_ fixation is observed through PEPc to facilitated starch synthesis. The activity of glycogen synthase (GS) is observed to vary according to previously mentioned pattern. Its activity is higher in MC than in GC at day, whereas, in the fourth phase, its activity is higher in GC than in MC (Figure 3a). During the daytime, starch synthesis in GC occurs from 3-phosphoglycerate through Calvin-Benson cycle (CBC) and plastidial fructose-1,6-bisphosphatase (pFBPase). However, during the night the conversion of 3-phosphoglycerate to hexose phosphate, a necessary step for starch production is halted when both of CBC and pFBPase are inactive. To facilitate starch synthesis glucose-6-phosphate directly enters the plastid to provide carbon skeletons in the 1^st^ phase of night (Figure 4a). This observation is also reported in previous experimental results (Daloso *et al*., 2017). The pattern of starch synthesis is also supported by the activities of different isoforms of PPDK in different phases. In phase 2 and 4, plastidial and cytosolic isoforms are active respectively, with higher activity in phase 4 than in phase 2 (Figure 3a,b). This supports the idea that gluconeogenesis in GC is enhanced at night to facilitate starch production at the beginning of the night (Daloso *et al*., 2016a). This starch is utilized mostly at night to provide carbon skeleton for glycolysis and mitochondrial metabolism to generate ATP. The rest of the starch is stored overnight and utilized in dawn (Horrer *et al*., 2016) to provide PEP for CO_2_ fixation by PEPc and malate for generating ATP through the TCA cycle. A higher activity of NADP-malate dehydrogenase is observed in chloroplast of GC than in MC at daytime in C_3_ (Figure 3a). This enzyme is responsible for interconversion of malate and OAA. OAA produced by PEPc at dawn is transported from cytosol to chloroplast. This OAA is reduced into malate by the enzyme NADP-MDH and the reducing power NADPH is supplied from the plastidial electron transport chain (pETC) in GC. Whereas in phase 2 and 3, there is no carbon fixation observed through PEPc and the OAA in chloroplast is entirely supplied from the mitochondria. However, in another aspect the NADP produced as a by-product by NADP-MDH is utilized in pETC to produce ATP, which is reported to be additional source of energy besides the ATP produced by mitochondrial electron transport chain (mETC) and glycolysis (Daloso *et al*., 2017). The redox in chloroplast is balanced in this way. The higher activity of NADP-MDH in GC than in MC is also experimentally reported in previous studies (Gotow et al., 1985; Shimazaki, 1989).

Whereas in CAM GC (Figure 2b), the starch production increases approximately twice of that in C_3_ GC. Here, starch synthesis begins at dawn, with the maximum starch synthesized during the first two phases of the day. In phase 1 and 2, the triose sugar produced by RuBisCO and the stored sucrose contribute in producing starch. Whereas in phase 3, starch production depends on the triose sugar produced in CBC. The stored starch begins to degrade in the first phase of the night, with the maximum amount degraded during the midnight. There is no starch storage at night, while the sucrose stored to maintain OP at night is used to produce starch at dawn. Amount of dark CO_2_ fixation by PEPc in GC is increased 3 times in CAM compared to C_3_. PEPc refixes a significant amount of respiratory CO_2_ into OAA, which finally gets converted to malate by MDH. Therefore, higher activities of PEPc and MDH are observed in CAM GC compared to C_3_ GC. However, this malate is stored and decarboxylated at daytime by ME only. Unlike C_3_ GC, there is no CO_2_ fixation observed by PEPc. In CAM GC maximum activities of plastidial and cytosolic PPDK are observed at daytime and G6P also enters plastid at day time. This supports the higher production of starch at daytime in CAM GC than in C_3_ GC. Stored starch is used to produce malate and citrate through the TCA cycle required for generating reductants in mitochondria to produce mitochondrial ATP at night and provide CO_2_ at daytime. However, citrate is stored approximately 10 times than malate and plays a major role in CO_2_ fixation in GC. In CAM GC, malate is itself stored at night and used in the first phase of the day, i.e., dawn. Hence, in CAM, there is no need for starch accumulation at night in GC.

However, in C_3_ MC (Figure 2a,3a,b), a minimal starch is produced and stored at daytime to supply carbon skeletons during the night for the production of phloem sap components and to generate ATP through glycolysis and the mETC. Malate is stored in a little amount in MC. Besides a little nocturnal accumulation of citrate is also observed in MC to provide carbon skeleton for the conversion to 2-oxogluterate at daytime. This pattern closely corresponds to experimental findings regarding C_3_ metabolism (Winter and Smith, 2022).

In CAM MC (Figure 2b), the starch production begins at dawn and continue throughout the day with the maximum production at midday. Here, starch production increases approximately 19 times than in C_3_ MC and higher activity of cytosolic PPDK is observed (Figure 4b). CO_2_ uptake in CAM MC is notably higher in the last phase of the night compared to the first two phases of the night. In fact, it is approximately 4 times the total CO_2_ uptake in the previous 11 hours. This CO_2_ is carboxylated into OAA through PEPc and converted to malate through MDH in cytosol. This malate is stored at night and gets decarboxylated at daytime by both PEPCK and ME, with approximately 14 times higher PEPCK activity than ME. To meet the increased demand of PEP, substrate of PEPc, during the night, a greater amount of starch is synthesized during the day. A little amount of citrate storage is also observed. At night, PEPc is active in CAM MC, but not in C_3_ MC. During the day, CO_2_ fixation is higher through RuBisCO than PEPc.

Unlike C_3_, in CAM, the activity of PPDK is observed in both MC and GC. At daytime, while both the plastidial and cytosolic forms are active in GC (Figure 4b), only cytosolic form is active in MC. At night, only a negligible activity of cytosolic PPDK is observed in GC. While the activities of mitochondrial enzymes like SDH, fumarase, and MDH are higher in MC at day, these are greater in GC at night.

### 2.5 Differential activities of C_4_ marker reactions in GC and MC

In C_3_ GC, the activity of PEPc during both day and night indicates a dark CO_2_ fixation alongside daytime CO_2_ fixation by RuBisCO and PEPc. Whereas in C_3_ MC, there is no nighttime CO_2_ fixation and CO_2_ is fixed only at daytime, by RuBisCO and PEPc with a very small activity of PEPc (Figure 3a). A higher activity of PEPc is observed in GC compared to MC during both light and dark phases (Figure 3a,b).

Furthermore, the activity of malate dehydrogenase in the chloroplast, cytosol, and mitochondria is observed to be higher in GC than MC (Figure 3a,b). The malate stored in the first phase to balance the charge of K^+^ is decarboxylated in the second phase by ME to provide CO_2_ for fixation during photosynthesis as there is maximum need for CO_2_ in chloroplast in this phase, with no activity of ME observed in MC. Microarray-based transcriptomic studies have confirmed these findings, showing elevated activities of enzymes which acts as C_4_ markers such as PEPc, ME, and MDH (described in previous section) etc., in GC. Taken together, these results suggest that GCs in C_3_ plants show metabolic characteristics related to C_4_ and CAM plants.

However, in CAM GC, RuBisCO is observed to fix the CO_2_ and facilitate starch synthesis at daytime. Whereas, at night CO_2_ is fixed by PEPc and stored as malate and citrate to provide carbon skeletons for CO_2_ production during the day. ME is active throughout all phases of the day to supply CO_2_ from this stored malate. The activity of PEPc in GC is higher in CAM than in C_3_. This result is in accordance with a previous experimental report on CAM GC, which showed that PEPc activity was up-regulated in GC when the metabolism shifted from C_3_ to CAM due to salt stress (Kong *et al*., 2020). In contrast, in MC of CAM, the activity of PEPc is higher than that in GC at night to fix the large amount of CO_2_ taken from the environment, into malate, which is stored overnight and decarboxylated by both phosphoenolpyruvate carboxykinase (PEPCK) and ME to provide CO_2_ for photosynthesis through RuBisCO. The flux through PEPCK is approximately 14 times higher than that through ME in MC of CAM. Our results indicate that the activities of PEPc and ME in GC are higher in CAM than in C_3_ plants, consistent with a previous experimental study by Kong et al., 2020.

The activity of RuBisCO is lower in GC compared to MC in both C_3_ and CAM. In C_3_ the activity of RuBisCO in GC is only 13% of that in MC, whereas in CAM it is 9%. This is due to the production of both the sucrose, transferred to GC and the phloem sap in MC. This suggests a lower photosynthetic capacity in GC compared to MC as previously experimentalized.

### 2.6 ATP production and utilization in GC and MC of C_3_ and CAM

Our findings suggest that in C_3_ plants, the ATPase proton pump in GC is primarily active during daytime when stomata are open. The overall ATP production during the daytime is greater in MC compared to GC (Figure 5a) because MC needs to produce starch at daytime for nighttime metabolic processes and phloem sap. Conversely, mitochondrial ATP production is higher in GC than in MC at daytime to support gluconeogenesis and meet the energy demand for osmolyte accumulation and driving the ATPase proton pump to open stomata (Figure 5a). Higher activity of mitochondrial MDH is also observed in GC than in MC during day (Figure 3a). Previous studies have highlighted the significance of ATP production via mitochondrial metabolism for stomatal opening (Daloso *et al*., 2017). At night, higher glycolytic and mitochondrial ATP production (Figure 5c) and higher activities of fumarase, SDH, and MDH are observed in GC compared to MC (Figure 3b), with greater CO_2_ production at night in GC. Greater O_2_ uptake in also observed in GC compared to MC, which is also reported in previous experiment (Shimazaki, 1989). These findings align with experimental results indicating higher activity of mitochondrial enzymes and respiratory rates in GC compared to MC (Mawson, 1993; Araújo *et al*., 2011; Vani and Raghavendra, 1994).

Conversely, in CAM plants, the ATPase proton pump activity is predominantly observed during the night. During the day, activities of mitochondrial fumarase, SDH, MDH are higher in MC than in GC, while at night, the reverse is observed. This nocturnal increase in mitochondrial metabolism supports its significance in facilitating stomatal opening. In MC, the ATP production through mETC and glycolysis is increased in the last phase of night to support the fixation of larger CO_2_ into malate. The enzyme PEPCK, necessary for the decarboxylation of stored malate, is ATP-dependent. The higher starch production and PEPCK activity are supported by increased ATP production through both pETC and mETC at daytime (Figure 5b,d).

Moreover, in MC, the ATP demand in CAM at day time is approximately 1.5 times higher than in C_3_, while at night it is approximately 6.4 times. This higher ATP production in CAM is required to meet the ATP demand to produce PEP from starch through glycolysis for CO_2_ fixation at night by PEPc. In C_3_, the ATP at day is utilized for for maintenance and gluconeogenesis pathway and at night utilized to supply maintenance cost. Whereas in CAM the requirement for ATP at night is for stomatal opening and CO_2_ assimilation and ATP is required at day for decarboxylation of stored malate. The similar pattern is observed in Shameer et al., 2018. However, in GC the ATP demand in C_3_ and CAM is approximately equal during both day and night. At night, the maximum ATP is utilized for maintenance and at day time besides providing maintenance cost, ATP is also utilized in gluconeogenesis process (Figure 5a,b,c,d).

Moreover, we observe different modes and flux through different reactions of TCA cycle to meet different demands of reductants for ATP production in GC and MC of C_3_ and CAM during day and night (Figure 4c,d,e,f). Notably, the full cyclic TCA cycle is active at night only. Flux through MDH and isocitrate dehydrogenase (IDH) is higher at day in GC in both C_3_ and CAM, whereas the flux through MDH increased tremendously in MC of CAM to provide high ATP at daytime.

Thus, in C_3_ plants, MC primarily focus on starch production during the daytime to serve as a source of food and to provide carbon skeleton for TCA cycle and mETC to produce ATP during the night. On the other hand, the metabolism in GC predominantly favours the activation of glycolysis and mitochondrial metabolism.

### 2.7 Metabolisms and energetics of GC and MC in partial CAM models like CAM cycling and CAM Idling

The distributions of flux of partial CAM models are given in Data S2. In CAM cycling, stomata are closed only at night. In MC, the CO_2_ generated by glycolysis and the TCA cycle, can’t leave the cell and is reconverted into malate by the enzyme PEPc at night. Subsequently, this malate is stored at night and decarboxylated during the day through PEPCK and ME. Both the anaplerotic CO_2_ and environmental CO_2_ taken by the MC are fixed by RuBisCO, leading to starch production from the triose sugars produced by RuBisCO. Whereas in the case of GC, starch production during the day is significantly increased to provide precursors of glycolysis and the TCA cycle for generating the required amount of ATP at night. The CO_2_ generated in these processes cannot leave the GC at night and is therefore converted into malate via PEPc, producing OAA. This OAA is then converted into citrate, which is also stored at night and decarboxylated during the daytime to release CO_2_. In GC of CAM cycling, the sucrose stored during the daytime to balance the OP at day is utilized in glycolysis at night. Proton-ATPase pumps send protons out of GC during the early morning and daytime when stomata are open, allowing K^+^ to enter in the morning to maintain OP.

In another variant of partial CAM known as CAM idling, both the GC and MC cease exchanging gases with the environment during both day and night. In MC no phloem sap is produced throughout the diel cycle. Both cell types function as isolated cells, with metabolism occurring in a closed manner. Unlike C_3_, CAM and CAM cycling, the energy demand of GC is higher compared to MC in CAM idling. In MC, daytime CO_2_ fixed by RuBisCO originates entirely from the anaplerotic CO_2_ derived from malate stored during the night. The resulting RuBisCO product is used to synthesize starch, which in turn fuels nighttime ATP production through respiration. The CO_2_ generated by respiration is then reconverted into malate via PEPc, thereby maintaining a closed cycle. On the other hand, in GC, the CO_2_ produced during respiration is fixed into citrate. Citrate is produced in mitochondria through citrate synthase and then stored until daytime, where it is decarboxylated to release CO_2_. In CAM Idling, there is no movement of protons and K^+^ into or out of the cell.

Results show that the photon requirement of MC is greater in CAM than C_3_, whereas in CAM cycling, it lies between C_3_ and CAM (Table S2). However, in case of CAM idling it is very low as there is no production of phloem sap. This pattern matches perfectly with Tay et al., 2021, whereas the photon demands of GC in C_3_, CAM and partial CAMs do not resemble this pattern. Unlike C_3_, CAM and CAM cycling, the photon demand of GC is higher compared to MC in CAM idling because the energy requirement in MC decreases without phloem sap production or sucrose transfer, whereas GC’s energy cost increases as it must cover its own maintenance expenses in this state.

The malate storage in MC during CAM cycling and idling are approximately 16% of that in CAM and the citrate storage in GC in both is increased to 6 times that in CAM. Unlike in MC, where malate is the primary stored metabolite at night, in GC, citrate becomes the predominant stored metabolite. There is no experimental report on the role of citrate storage in GC. We may get this due to the constraints in our modelling. However, experimental studies are further needed to verify the role of citrate in GC.

### 2.8 Analyses by varying the constraints

Since there are no experimental reports about the amount of sucrose transferred from MC to GC, we explored various percentages of sucrose entering GC and simulated the model in C_3_ condition (Data S3). While the percentage of sucrose transfer increases, the photon demand of GC declines as it starts fulfilling its energy requirements from the sucrose influx. Conversely, the photon demand of the MC escalates with increased sucrose transfer, as it must generate the sucrose. Similar patterns are observed in RuBisCO’s carboxylase activity and mETC and pETC. Moreover, as sucrose gets converted into starch for nighttime utilization in GC, starch production increases with a higher influx of sucrose. Relative population of GC and MC may control how much sucrose per MC will be available to transport to one GC. The consistency of observed metabolic pattern at varying percentage of transported sucrose indicate the model is good enough to predict some important aspects of metabolism.

We further varied the ratio of length of day and night and observed that in general, in long day conditions (day:night = 16:8 and 14:10), the photon demands of both the cells decrease due to lower starch production for shorter dark phase, whereas in short day conditions (day:night = 8:16 and 10:14), the photon demands of both the cells increase because of higher starch need at dark phase in all the four variants (Table S3). Moreover, the photon demand of MC is greater than that of GC in all the conditions.

Besides analysing the results in the standard scenario, we have simulated our model using only Cl^-^ as counterion of K^+^ (Distributions of flux for all the four metabolic variants are given in Data S4.) and compared the results with the standard scenario and observed that accumulation of these ions are energetically similar (Table S4). However, the activities of the pathways and enzymes related to malate production and utilization in GC show major differences in these two scenarios (Listed in Table S5). The total starch synthesis is higher in standard scenario. The synthesis of starch begins at mid-day, peaks in the 1^st^ phase of night and the produced starch is stored at night to provide the precursor for malate synthesis at dawn. On the other hand, when only Cl^-^ acts as counterion (i.e., there is no accumulation of malate at daytime), enzymes related to metabolism of malate like PEPc, ME, PPDK etc., are inactive at day time and starch is synthesized only in the 1^st^ phase of night. The decreased starch production and no night time storage of starch in this scenario are due to zero demand of malate at daytime. RuBisCO activity in phase 1 and 2 also varies in these two scenarios depending on the different patterns of starch production. Previous study on isolated C_3_ GC (Tan and Cheung, 2020) observed similar results using only Cl^-^ as counterion. However, in addition, even when only Cl^-^ is used as counterion instead of malate, our model is able to predict the higher activities of some of the C_4_ like enzymes like NADP-MDH, fumarase, SDH, MDH etc., in GC than in MC of C_3_ as experimentally reported (Daloso *et al*., 2017). We have further checked for the preferred counterion of K^+^ in the model allowing both malate and Cl^-^ to accumulate in GC. The model prefers only Cl^-^ to accumulate. However, experimental studies reported that malate acts as a major counterion to balance the charge of K^+^ (Lee, 2010). This suggests that there may be some additional constraints due to which GC uses malate as major counterion, which could not be captured in the model (when both malate and Cl^-^ are to accumulate in GC). Moreover, we have also analysed the model using the dual optimization criteria of maximizing the phloem sap production followed by the minimization of the total cellular flux and observed no differences.

### 2.9 Scopes and future aspects

The six phase combined diel model of GC and MC successfully explains many experimental observations and computational predictions. This combined model would help to integrate and model metabolism of all the cells present in a leaf and to explain physiological process starting from gaseous exchange to biosynthesis and transfer of photosynthates to phloem sap (Hunt *et al*., 2023). The inclusion of malate transport from MC to GC (Lee, 2010; Lawson *et al*., 2014, Daloso *et al*., 2017), population based study of GC and MC considering the different numbers of GCs and MCs in a leaf, integration of omics data, dependencies on CO_2_ concentration, restriction on vacuolar size, regulation of hormones like abscisic acid (ABA), roles of calcium signalling etc. (Lawson, 2009; Lawson *et al*., 2014), will help us to get more detailed and realistic accounts of the leaf metabolism.

## 3. Experimental procedures

### 3.1 Constraints applied to simulate guard cell and mesophyll cell

A well-established relation of osmotic pressure, stomatal opening, and gaseous exchange is mimicked in our model. When osmolytes like KCl, K_2_Malate, sucrose etc., accumulate in a phase, the OP rises in GC and the CO_2_ can enter the internal airspace of the leaf. Both GC and MC are linked to internal airspace. Exchange of O_2_ between the internal airspace and environment also depends on stomatal opening and closing and thus on OP in such a way that as the OP increases, the upper limit of O_2_ exchange escalates. Initially, the combined model is simulated under the constraints representative of a C_3_ plant. In C_3_, the OP in GC in the three phases of day is assumed to be 1.39 µosmol.m^-2^.s^-1^ (Methods S1). This is equivalent to 900 femto-osmol/cell/h (Reckmann et al., 1990) considering the average area of GC as 180 µm^2^ (Melaragno *et al*., 1993). Values of osmotic coefficients of KCl, K_2_Malate and sucrose are used from Tan and Cheung, 2020. To simulate CAM and partial CAMs like CAM cycling and CAM idling we constrain the gaseous exchange of internal air-space with the cells (Table S6). In CAM plants, previous experimental reports suggest that stomata are entirely closed only at midday. Therefore, we block gaseous exchange completely in phase 2 and set the OP to 0. To maintain a partial gaseous-exchange as experimentally observed in 1^st^ and 3^rd^ phase of day (Heyduk, 2022), we set the OP to 0.2 unit. In all the phases of night OP is 1.39 unit. For CAM cycling, the OP is applied in the same manner as C_3_ and the gaseous exchange is blocked at night, whereas in CAM idling, OP is set to 0 in all the phases and there is no gaseous exchange. Maintenance cost of GC is calculated following Tan and Cheung, 2020 and that for MC are utilized from Tay et al., 2021. All the other constraints are listed in the Table 1.

Experimental studies have highlighted the significant roles of metabolites such as sucrose, synthesized in MC, in regulating stomatal movement. Sucrose holds particular importance as an osmolyte in GC, with experimental evidence suggesting its transport from MC to GC during stomatal opening (Lee, 2010; Lawson *et al*., 2014). We also allow the sucrose transfer from MC to GC during the stomatal opening. The amount of sucrose transported is set at 80% of the total osmolytes required to maintain the OP in GC as in Tan and Cheung, 2020.

As there is no report on accumulation of osmolytes at night in C_3_, we have imposed a constraint to restrict the transfer of K^+^ from day to night in our simulation. However, no such constraint is applied on K^+^ storage in CAM plants, as K^+^ accumulation is necessary to maintain OP in night as well as day. Additionally, research suggests that both malate and Cl^-^ can serve as counterions for K^+^. However, Lee, 2010 reported that malate plays a more significant role to balance the positive charge of K^+^. We have analysed our results using both malate and Cl^-^ separately as counterions for K^+^. The Table 1 enlists the general constraints used in GC-MC metabolisms in all the four variants.

### 3.2 Flux Balance Analysis

Flux Balance Analysis (FBA), a constraint-based modelling approach is used here for simulating our model. In general, the objective used in this study is to minimize the total absolute sum of fluxes (proxy of minimization of cellular economy), called parsimonious flux balance analysis (pFBA). Further, we have analysed our model using a dual optimization-maximization of phloem sap production, followed by minimization of the total absolute sum of fluxes. In case of dual optimization, we have used the value of photon obtained from the previous pFBA result as constraint. The scobra package (Shaw and Cheung, 2018) is used for simulations of FBA. The details of model construction and methodology are presented in Methods S1 and S2 and the codes for simulation are given in Data S5.

## Acknowledgements

DS thanks Department of Biotechnology, Government of India for her fellowship.

## 5. Author contributions

DS and SK designed the study, DS performed the simulations, both interpreted the results and co-wrote the paper.

## 6. Data availability

The model (in sbml and excel format) and the codes are provided in Data S1 and S5 respectively.

## 7. Supporting information

**Table S1** Comparison of our study with two of the previous studies.

**Table S2** Energetic differences in C_3_, CAM, and partial CAMs.

**Table S3** Different photon demands of GC and MC of all four variants in different photoperiods.

**Table S4** Photon demands of GC and MC of the four metabolic variants in two scenarios using Cl^-^ and malate^2-^ separately as counterions of K^+^.

**Table S5** Different enzymatic activities in C_3_ GC using malate^2-^ and Cl^-^ separately as counterion of K^+^.

**Table S6** Different gaseous constraints used in our simulations.

**Methods S1** Detailed calculation of osmotic pressure (OP).

**Methods S2** Details of Flux Balance analysis (FBA).

**Data S1** Six phase combined model of guard cell and mesophyll cell in sbml and excel formats.

**Data S2** Flux solutions for C_3_, CAM, CAM cycling and CAM idling conditions in standard scenario.

**Data S3** Flux solutions for different percentages of sucrose transfer from MC to GC.

**Data S4** Flux solutions for C_3_, CAM, CAM cycling and CAM idling conditions using only Cl^-^ as the counterion of K^+^.

**Data S5** Codes for simulations of the model in all the four variants and sucrose scan.

