## Supporting information for "Lessons from combined metabolic model of mesophyll and guard cells"

The following Supporting Information is available for this article:

**Table S1** Comparison of our study with two of the previous studies.

**Table S2** Energetic differences in C_3_, CAM, and partial CAMs.

**Table S3** Different photon demands of GC and MC of all four variants in different photoperiods.

**Table S4** Photon demands of GC and MC of the four metabolic variants in two scenarios using Cl^-^ and malate^2-^ separately as counterions of K^+^.

**Table S5** Different enzymatic activities in C_3_ GC using malate^2-^ and Cl^-^ separately as counterion of K^+^.

**Table S6** Different gaseous constraints used in our simulations.

**Methods S1** Detailed calculation of osmotic pressure (OP).

**Methods S2** Details of Flux Balance analysis (FBA).

**Data S1** Six phase combined model of guard cell and mesophyll cell in sbml and excel formats.

**Data S2** Flux solutions for C_3_, CAM, CAM cycling and CAM idling conditions in standard scenario.

**Data S3** Flux solutions for different percentages of sucrose transfer from MC to GC.

**Data S4** Flux solutions for C_3_, CAM, CAM cycling and CAM idling conditions using only Cl^-^ as the counterion of K^+^.

**Data S5** Codes for simulations of the model in all the four variants and sucrose scan.

**Table S1** The table enlists the experimentally observed trends of some important metabolic enzymes, osmolyte accumulation and respiratory rate in GC and MC that have not been observed in previous studies. In 2017, Robaina-Estévez *et al.*, 2017 examined metabolic flux variations between MC and GC in *Arabidopsis thaliana*. Meanwhile, a study on a four phase metabolic model of isolated C_3_ guard cells delved into the metabolism, illustrating their role in balancing charge and accumulating osmolytes to regulate OP (Tan and Cheung, 2020). These investigations provide valuable insights into the distinct metabolic processes of GC. Whereas, our six-phase combined model integrates both guard cells (GC) and mesophyll cells (MC), offering a more realistic portrayal of metabolic dynamics. This model specifically highlights metabolic differences and varying enzyme activity levels within GC and MC across C_3_, CAM, and partial CAM metabolisms. By focusing on these differences, our model accurately predicts outcomes, aligning closely with experimental studies in C_3_ metabolism (Table 2 in main text). Furthermore, it hypothesizes the interconnected metabolism in CAM and partial CAMs, providing valuable insights into these complex metabolic pathways.

| Our combined model of GC and MC | Robaina-Estévez *et al*., 2017 | Tan & Cheung, 2020 |
| --- | --- | --- |
| Osmolyte accumulation in CAM in different phases | NO | NO |
| PEPc activity at night in C_3_ | NO | NO |
| Higher activity of NADP-MDH in C_3_ GC | YES | NO |
| Lower RuBisCO activity in GC than MC in both C_3_ and CAM | NO | NO |
| Different activities of ME in different phases in GC and MC of C_3_ and CAM | Only in light | Only in GC |
| Activity of PPDK in different phases of GC and MC of C_3_ and CAM | NO | NO |
| Activity of fumarase, SDH in different phases of GC and MC of C_3_ and CAM | NO | NO |
| pFBPase and PPDK activity at night in C_3_ GC | NO | NO |
| Higher respiration in GC than in MC in both C_3_ and CAM | NO | NO |

**Table S2** The table contains the photon demands and fluxes (Unit: µmol.m^-2^.s^-1^) through mitochondrial and plastidial ATP productions in C_3_, CAM and partial CAMs during both day and night. In different metabolic conditions the photon demand and ATP production through mitochondrial electron transport chain (mETC) and plastidial electron transport chain (pETC) varied in GC and MC during both day and night. For MC, the photon demand of CAM is greater than that in C_3_, whereas for CAM cycling it lies between the photon cost of C_3_ and CAM. However, CAM idling requires least amount of photon as there is no production of phloem sap by MC. Moreover, result shows that the photon demands of GC does not match this pattern. For C_3_ and CAM, the photon demand does not vary, whether it increases for partials CAM and becomes maximum for idling as there is no sucrose transfer in CAM idling and the GC has to produce metabolites for its own maintenance throughout the day and night.

|  | *C_3_* | *CAM* | *CAM cycling* | *CAM Idling* |
| --- | --- | --- | --- | --- |
| *Photon in GC* | 153.90 | 146.27 | 204.82 | 343.70 |
| *Photon in MC* | 394.36 | 441.68 | 413.93 | 64.93 |
| *pETC in GC* | 16.49 | 15.67 | 21.94 | 36.82 |
| *pETC in MC* | 42.94 | 47.32 | 45.16 | 6.96 |
| *mETC at day in GC* | 10.88 | 11.92 | 20.41 | 21.96 |
| *mETC at night in GC* | 15.07 | 14.29 | 15.66 | 15.39 |
| *mETC at day in MC* | 5.53 | 24.45 | 7.41 | 4.41 |
| *mETC at night in MC* | 2.50 | 11.10 | 3.78 | 3.69 |

**Table S3** The table contains the different amounts (Unit: µmol.m^-2^.s^-1^) of required photon in different ratios of day and night in the four metabolic conditions. In the main text, we have given the results assuming the ratio of day and night to be 12:12. Changes of photon demands with changes in different ratios of day and night are listed below. Results show, as we decrease the length of night, the cells have to store less amount of starch for continuing night time metabolism and maintenance and the photon cost decreases, whereas when night time increases, the photon demand increases for the same reason in the both cells of all the four conditions.

| Ration of day and night | | 8:16 | 10:14 | 12:12 | 14:10 | 16:8 |
| --- | --- | --- | --- | --- | --- | --- |
| C_3_ | GC | 192.51 | 173.22 | 153.90 | 134.68 | 128.13 |
|  | MC | 390.36 | 396.02 | 394.36 | 390.11 | 385.43 |
| CAM | GC | 180.32 | 163.32 | 247.27 | 130.20 | 124.87 |
|  | MC | 446.46 | 442.98 | 441.68 | 439.92 | 437.40 |
| CAM cycling | GC | 269.19 | 233.57 | 204.82 | 176.12 | 161.70 |
|  | MC | 430.41 | 422.39 | 413.93 | 405.08 | 396.80 |
| CAM idling | GC | 388.06 | 365.87 | 343.70 | 321.51 | 299.33 |
|  | MC | 75.41 | 70.17 | 64.93 | 59.69 | 54.57 |

**Table S4** The table enlists the amount of photon (Unit: µmol.m^-2^.s^-1^) utilized by GC and MC in the four metabolic conditions in the two scenarios- i) in our standard scenario, where we have allowed only malate to act as a counterion of K^+^ and ii) another scenario, where we have allowed only Cl^-^ to act instead of malate and results show that both works equivalently in case of photon utilization. (* The values of photon demands in CAM idling do not change as there is no osmolyte accumulation observed in GC)

| Osmolyte used as counterion of K^+^ in GC | C_3_ | | CAM | | CAM cycling | | CAM idling^*^ | |
| --- | --- | --- | --- | --- | --- | --- | --- | --- |
|  | MC | GC | MC | GC | MC | GC | MC | GC |
| Mal only | 394.36 | 153.90 | 440.06 | 147.45 | 412.55 | 206.09 | 64.93 | 343.69 |
| Cl^-^ only | 394.36 | 154.20 | 440.06 | 147.69 | 412.55 | 205.39 | 64.93 | 343.69 |

**Table S5** The table enlists the differences of activities of enzymes observed when only malate and only Cl^-^ act as counterion to balance the positive charge of K^+^ (Unit: µmol.m^-2^.s^-1^).

|  | | Only malate^2-^ | Only Cl^-^ |
| --- | --- | --- | --- |
| Starch accumulation in GC | Phase 1 to phase 2 | 0 | 0 |
|  | Phase 2 to phase 3 | 0.20 | 0 |
|  | Phase 3 to phase 4 | 0.25 | 0 |
|  | Phase 4 to phase 5 | 2.23 | 1.98 |
|  | Phase 5 to phase 6 | 0.47 | 0.22 |
|  | Phase 6 to phase 1 | 0.25 | 0 |
| PEPc | Phase 1 | 0.58 | 0 |
|  | Phase 4 | 0.68 | 0.68 |
| ME | Phase 2 | 0.58 | 0 |
|  | Phase 5 | 0.68 | 0.68 |
| PPDK | Phase 2 | 0.58 | 0 |
|  | Phase 4 | 1.39 | 1.07 |
| RuBisCO | Phase 1 | 0.31 | 0.17 |
|  | Phase 2 | 3.34 | 3.38 |
|  | Phase 3 | 0.35 | 0.36 |

**Table S6** The table includes the constraints on gaseous exchange (CO_2_ and O_2_) in the four metabolic conditions.

| Constraints | C_3_ | CAM | Cycling | Idling |
| --- | --- | --- | --- | --- |
| CO_2_ exchange in MC at day | Unconstrained | Partly allowed | Unconstrained | 0 |
| CO_2_ exchange in MC at night | Unconstrained | Unconstrained | 0 | 0 |
| CO_2_ exchange in GC at day | Unconstrained | Partly allowed | Unconstrained | 0 |
| CO_2_ exchange in GC at night | Unconstrained | Unconstrained | 0 | 0 |
| O_2_ exchange in MC at day | Unconstrained | Partly allowed | Unconstrained | 0 |
| O_2_ exchange in MC at night | Unconstrained | Unconstrained | 0 | 0 |
| O_2_ exchange in GC at day | Unconstrained | Partly allowed | Unconstrained | 0 |
| O_2_ exchange in GC at night | Unconstrained | Unconstrained | 0 | 0 |

**Methods S1**

The rate of solute accumulation in an intact guard cell is reported to be 900 femto-osmol/cell/h (Reckmann *et al*., 1990) and the average length and width of guard cells are reported to be 20µm and 9µm (Melaragno *et al.*, 1993). From these to data, we have calculated the rate of solute accumulation per m^2^ per s.

$$\frac{900 femto-osmol}{cell*h}=\frac{900*{10}^{-9} \mu osmol}{180* {10}^{-12}m^{2}*3600s}=1.39 \mu osmol.m^{-2}.s^{-1}$$

**Methods S2**

Flux Balance Analysis (FBA) is a constraint-based modelling approach that allows the identification of optimal flux through the reactions of a metabolic network in a steady-state by maximizing or minimizing the objective function, defined according to the desired objective. Our models are simulated using FBA (Orth *et al.*, 2010). Different constraints were used for GC and MC and the objective function is minimization of total cellular flux. In our study, simulations of FBA has been done using the scobra package (Shaw & Cheung, 2018).

At steady state, the rate of production and the rate of consumption are equal for each internal metabolite in the model and that can be mathematically represented as:

**Sv** = 0

where, the vector **v** represents the flux through all the n number of reactions and **S** is the stoichiometry matrix of size m x n, where m represents the number of metabolites and n represents the number of reactions of the network.

The optimization problem can be defined as,

maximize or minimize **z** = **c***^T^***v**

The flux constraints are given as,

**c***_l_* ≤ **v** ≤ **c***_u_*

Where, **z** is the objective function, **c***^T^* is the transpose of a vector (**c**) of weights that indicate how much a reaction contributes to the objective, **v** is the vector of all fluxes, **S** is the stoichiometry matrix and **c***_l_* and **c***_u_* are the vectors of lower and upper bound of fluxes, respectively. In case of irreversible reaction, the lower bound **c***_l_* becomes 0 and allowable flux is limited to be greater than or equals to 0.

**Data S1, S2, S3, S4 and S5** are submitted as separate files during submission.
